# Adolescent stress-induced ventral hippocampus redox dysregulation underlies behavioral deficits and excitatory/inhibitory imbalance related to schizophrenia

**DOI:** 10.1101/2023.11.28.568850

**Authors:** Thamyris Santos-Silva, Caio Fábio Baeta Lopes, Doğukan Hazar Ülgen, Danielle Aparecida Guimarães, Francisco S. Guimarães, Luciane Carla Alberici, Carmen Sandi, Felipe V. Gomes

## Abstract

Redox dysregulation has been proposed as a convergent point of childhood trauma and the emergence of psychiatry disorders, such as schizophrenia (SCZ). However, the impact of severe stressors during adolescence on the ventral hippocampus (vHip) redox states and their functional consequences, including behavioral and electrophysiological changes related to SCZ, are not entirely understood. After exposing adolescent animals to physical stress (postnatal day, PND31–40), we explored social and cognitive behaviors (PND47–49) and the activity of pyramidal glutamate neurons, the number of parvalbumin (PV) interneurons, and the transcriptomic signature of the vHip (PND51). We also evaluated the impact of stress on the redox system one and ten days after stress, while glutathione levels were measured in the vHip and serum following the behavioral test. Adolescent-stressed animals exhibited loss of sociability, cognitive impairment, and vHip excitatory/inhibitory (E/I) imbalance. Genome-wide transcriptional profiling unveiled the impact of stress on synaptic and redox system-related genes. Stress impacted mitochondrial respiratory function, leading to changes in reactive oxygen species levels in the vHip. Glutathione (GSH) and glutathione disulfide (GSSG) levels were elevated in the serum of stressed animals, while GSSG was also increased in the vHip and negatively correlated with sociability. Additionally, PV interneuron deficits in the vHip caused by adolescent stress were associated with oxidative stress. Our results highlight the negative impact of adolescent stress on vHip redox regulation and mitochondrial function, which are partially associated with E/I imbalance and behavioral abnormalities related to SCZ.

## Introduction

Adverse events during neurodevelopmental phases, such as childhood and adolescence, have been identified as significant risk factors for the emergence of schizophrenia (SCZ)^1^. For instance, individuals who have experienced severe trauma during these periods are 3.5 times more likely to have psychosis in adulthood^2^. Notably, social and cognitive deficits are also symptoms of SCZ and precedes psychosis episodes, which usually occurs during late adolescence/early adulthood and may lead to the SCZ diagnosis^3^. Given that early adolescence is a sensitive period for experience-dependent plasticity and adverse environmental experiences impact behavioral outcomes later in life^4,5^, understanding the molecular mechanisms that govern social and cognitive dysfunction under severe early adolescent stress is a critical step to discerning the underpinnings of the emergence of SCZ.

During early adolescence, the GABAergic parvalbumin (PV) interneurons are highly plastic in terms of excitatory drive and neuronal activity, making them particularly susceptible to damage by stressors^6^. Following adolescence, extracellular matrix structures named perineuronal nets (PNNs) are formed around PV interneurons, protecting them from stress-induced damage mediated by oxidative stress^7–9^. Accordingly, our previous studies have indicated a causal role of adolescent stress and reduction in PV interneurons and their associated PNNs, increased pyramidal neuron activity in the ventral hippocampus (vHip), and behavioral changes in adulthood associated with SCZ^10^. Importantly, hippocampal hyperactivity is proposed to induce an overactive dopamine system^11^, which has been consistently associated with psychotic symptoms in individuals with SCZ and behavioral abnormalities in our animal model^10,12^.

Several studies have pointed to redox dysregulation as one “hub” associated with the emergence of psychosis in early adulthood^13^. A dysregulated stress response system leads to deficits in energy production and enhanced reactive oxygen species (ROS) generation, which overwhelms the antioxidant capacity and has been linked to oxidative stress and cell damage^14^. PV interneurons have elevated metabolic activity and oxidative phosphorylation to support their high-frequency activity, making them particularly vulnerable to oxidative stress and increased metabolic demand^15^. Most studies on oxidative stress-induced PV interneuron deficits focus on SCZ animal models utilizing prenatal/neonatal manipulations, genetic approaches, or insults during adulthood^16–22^. However, the relationship between environmental stress exposure during the postnatal period and long-term redox homeostasis, mitochondrial function, and behavioral and circuitry changes associated with SCZ have been less explored.

Here, we studied the behavioral impact of early adolescent stress exposure in rats and changes in the excitatory/inhibitory (E/I) balance by evaluating the number of PV-positive cells and pyramidal neuron activity in the vHip. After gaining insights by employing a transcriptomic analysis, we investigated redox regulation and the association of oxidative stress with behavioral abnormalities and co-localization with PV interneurons. Furthermore, we explored whether glutathione (GSH and GSSG forms) levels, a major intracellular antioxidant, are altered in the vHip and serum of stressed animals and their correlation with behavioral outcomes. Altogether, we gained insights into a putative mechanism underlying the impact of adolescent stress and the emergence of behavioral and neural circuit changes related to SCZ.

## Methods

A detailed description of all experimental procedures, including animals, stress protocol, behavioral tests, electrophysiology, transcriptomic analysis, mitochondrial respirometry, biomolecular assays, and immunofluorescence, is provided in the supplementary material.

### Animals

Male Sprague-Dawley rats were housed (2–3 animals per cage) in a temperature-(22 °C) and humidity-(47%) controlled environment (12 h light/dark cycle; lights on at 6 AM) with water and food *ad libitum*. All procedures were approved by the Ribeirão Preto Medical School Ethics Committee (155/2018 and 248/2019), which follows Brazilian and International regulations.

### Stress procedure

Animals were exposed to inescapable footshock (FS; from PND 31–40) daily and three restraint stress (RS) sessions (PD31, 32, and 40). Using this protocol, we previously found marked behavioral and electrophysiological changes resembling SCZ that persist until adulthood^10,23,24^.

### Behavioral tests

Naïve and stressed animals were subjected to social interaction and novel-object recognition tests from PND 47 until 50, as previously described^23,24^.

### In vivo recordings of vHip pyramidal neurons

Pyramidal neurons were identified by typical electrophysiological characteristics such as firing rate and action potential shape and registered as previously described^10,23^.

### Immunofluorescence

All immunofluorescence procedures, image acquisition, and analysis followed previously reported protocol^10^.

### Gene expression profiling from the rat vHip

At PND 51, RNA was isolated from vHip samples, and then bulk-RNA sequencing, transcriptomic analysis, and functional gene set enrichment analysis were performed as previously described^25^.

### MitoSOX™, AmplexRed® and Glutathione/Glutathione disulfide (GSH/GSSG) assay

To measure the release of superoxide production by mitochondria from vHip samples, we used the MitoSOX™ mitochondrial superoxide indicator (ThermoFisher, M36008) assay. Amplex® Red Hydrogen Peroxide/Peroxidase Assay Kit (ThermoFisher, A22188) was used to measure hydrogen peroxide and peroxidase activity. GSH and GSSG levels were measured by a green fluorescence assay kit (Abcam, ab205811).

### High-resolution respirometry

2 mg of fresh vHip samples from both hemispheres, cut into fine fragments, were properly collected to measure mitochondrial respiration using an Oxygraph-2k respirometer (Oroboros, Austria), as previously described^26^.

### Statistical analyses

All data were subjected to tests to verify the homogeneity of variances (Bartlett’s test) and if they followed a normal distribution (Shapiro–Wilk test), and then properly subject to parametric or non-parametric analyses. Significant differences were indicated by p < 0.05.

## Results

### Adolescent stress exposure causes behavioral deficits and affects inhibitory and excitatory neurons in the vHip

Our previous findings emphasize the importance of adolescence as a critical period of vulnerability and the emergence of SCZ-like symptoms in adulthood^10,24^; however, it was unknown if behavioral changes were already present in early periods, i.e., during late adolescence. Hence, we evaluated the effect of adolescent stress on sociability and cognitive function during late adolescence (PND47–50), and then assessed the E/I balance in the vHip (PND51–63) (Figure 1A). Stressed animals showed reduced social interaction time (SIT) with an unfamiliar rat (Figure 1B) and less novel object recognition index (NOR), as revealed by the discrimination index (Figure 1C), demonstrating the harmful impact of adolescent stress on social and cognitive performance that appeared one week after stress exposure.

**Figure 1.**
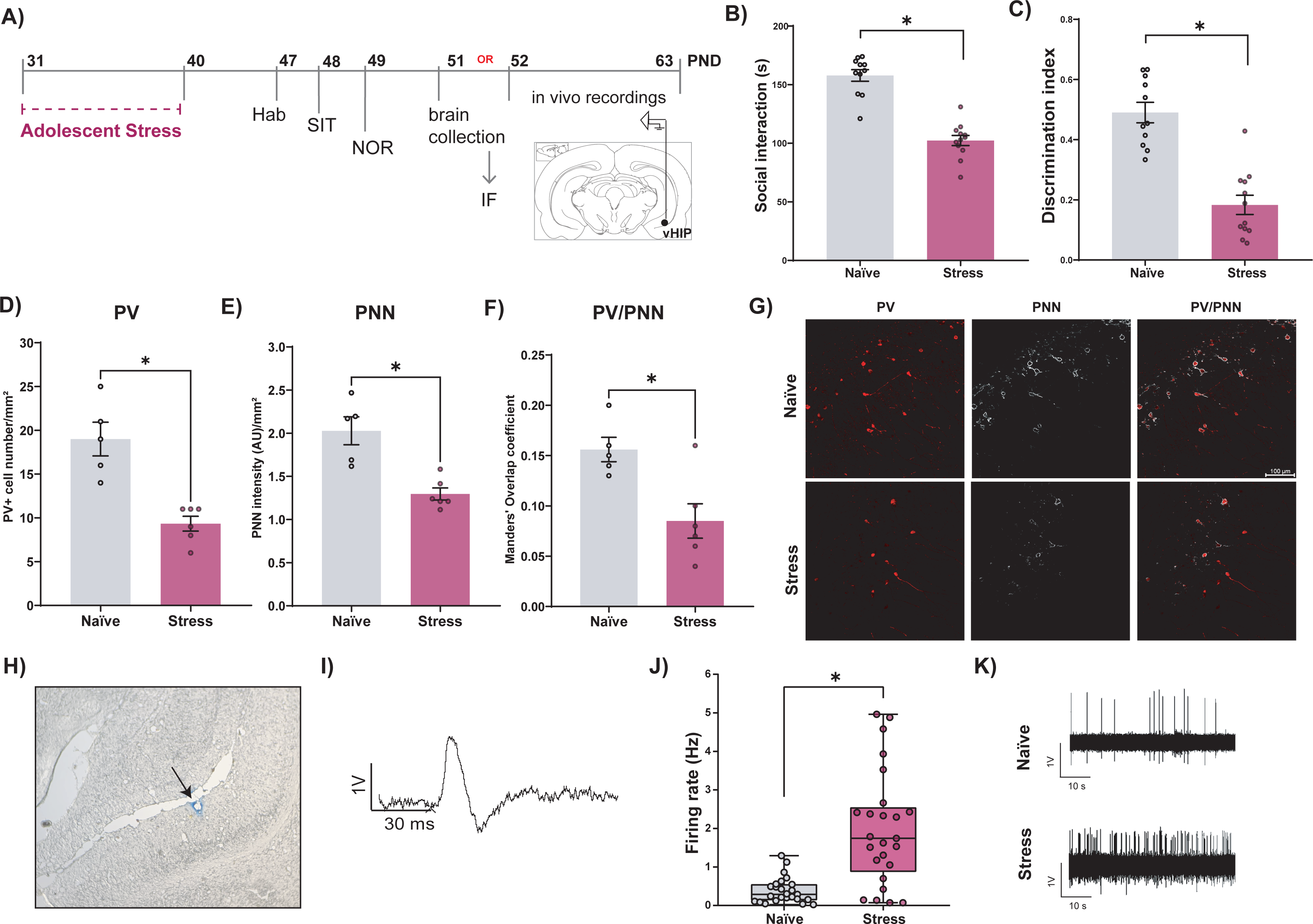
Adolescent stress leads to social and cognitive deficits and impacts PV interneurons, their associated PNNs, and vHip pyramidal neuron activity. **(A)** Experimental design. **(B)** Adolescent stress decreased social interaction time (t_21_ = 8.46, p < 0.0001), and **(C)** impacted the discrimination index in the NOR test (t_21_ = 6.57, p < 0.0001) (n = 11-12/group). **(D)** Adolescent stress reduced the number of PV positive cells/mm² (t_9_ = 4.90, p = 0.008), **(E)** PNN fluorescence intensity/mm² (t_9_ = 4.43, p = 0.002), and **(F)** PV/PNN co-localization (t_9_ = 2.67, p = 0.01). Data are shown as mean ± SEM. *p < 0.05; unpaired t-test. **(G)** Representative image of immunohistochemical staining of parvalbumin (red), Wisteria floribunda agglutin-labeled PNN (write) and PV and PNNs co-localization in the rat ventral hippocampus of naïve and stressed animals. **(H)** The black arrow indicates the final location of the last track that was marked by electrophoretic ejection of dye for histological verification. **(I)** Typical electrophysiological characteristics of pyramidal neurons in the vHip. **(J)** Adolescent stress increased the firing rate of pyramidal neurons in the vHip 1–2 weeks after stress (naive group: n = 25 neurons from 6 rats, 0.39 ± 0.06 firing rate; stress group: n = 25 neurons from 6 rats, 1.97 ± 0.30 firing rate). Data are represented as boxes and whiskers (minimum and maximum values). U = 99; *p < 0.05; Mann-Whitney test. **(K)** Representative traces of spike activity from vHip pyramidal neurons from naïve and stressed animals.

PV interneurons are highly vulnerable to stress during adolescence, and the formation of their associated PNNs is not complete until early adulthood^1,27^. Stress exposure during adolescence decreased the number of PV-positive cells at PND51 (Figure 1D) and the fluorescence intensity of PNNs (Figure 1E). It also impacted the Mander’s overlap coefficient of PV-positive cells and PNNs fluorescence, indicating a decreased PV and PNN co-localization (Figure 1F). Representative images are shown in Figure 1G. Given that early adolescent stress reduced the number of PV-positive cells, which could change the E/I balance, we recorded the activity of glutamatergic pyramidal neurons in the vHip (PND 52–63; Figure 1H and I). Increased firing rate of vHip pyramidal neurons was observed in stressed animals (Figure 1J). Representative traces of spike activity from vHip pyramidal neurons from naïve and stressed animals are shown in Figure 1K. These findings indicate that adolescent stress led to vHip E/I dysregulation.

### Transcriptomic analysis of the vHip reveals changes in synaptic and redox regulation-related genes

Next, we performed a hypothesis-free transcriptomic analysis to explore the effects of adolescent stress on vHip (see Supplementary Table 1 for a list of differentially expressed genes (DEGs) in the vHip). Considering DEGs p-value < 0.01, we found 57 down-regulated and 54 up-regulated in the vHip of stressed animals (Figure 2A). Applying a gene set enrichment analysis (Supplementary Table 2), we identify significant pathways associated with the redox system, such as “Negative Regulation Of Oxidative-stress Induced Neuron Death” and “Glutathione Transferase Activity” (Figure 2B). Notably, stress also enriched pathways related to “Negative Regulation Of Intrinsic Apoptotic Signaling Pathway In Response To DNA Damage”. Moreover, synaptic gene ontology (SynGO) analysis, an expert-curated resource for synapse function gene enrichment studies, revealed significantly enriched biological processes related to “Regulation Of Synaptic Vesicle Cycle” (Figure 2C). Those findings suggest that E/I imbalance is associated with changes in the presynaptic neurons, possibly due to loss of extracellular matrix around PV interneurons and oxidative damage in neuronal cells. Indeed, we found an enrichment in “Ing L5-6 PVALB MEPE - BRAIN” (p-value = 0.03) by analyzing DEGs on the Human BioMolecular Atlas Program (HuBMAP ASCTplusB augmented 2022), suggesting that enriched DEGs are more likely to be found in the PV interneurons (Supplementary table 2).

**Figure 2.**
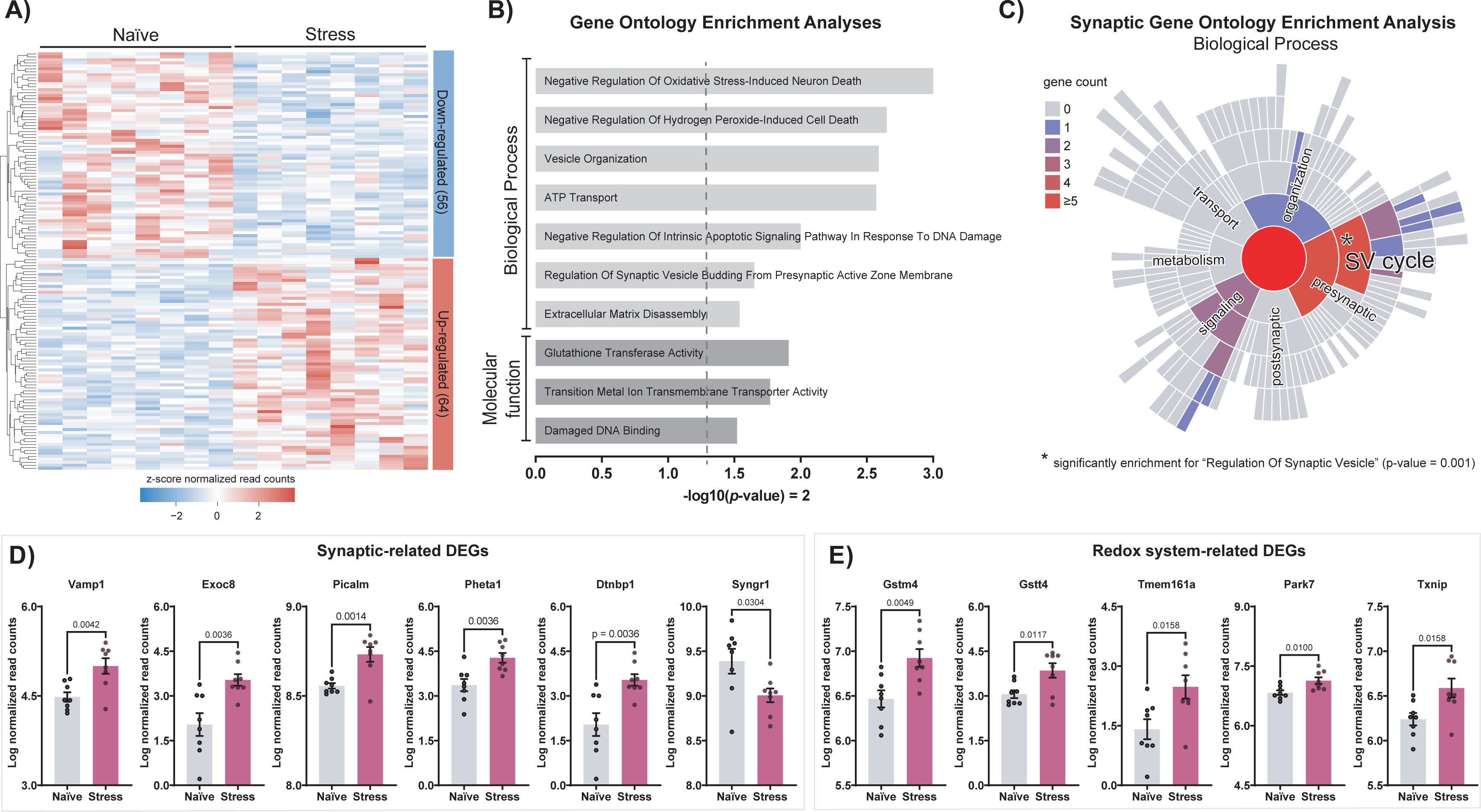
Gene expression changes in the vHip of stressed animals. **(A)** Heatmap of normalized mRNA expression levels of differentially expressed genes (DEGs, p-value < 0.01) in the vHip comparing adolescent stress vs. naïve groups. The overall numbers of DEGs up and down-regulated are described in their respective columns. The dendrograms represent the hierarchical clustering of genes according to gene expression (Euclidean distance method). **(B)** Select gene set enrichment analyses of biological process and molecular function ontologies (p-value < 0.05) considering DEGs in the vHip of stressed animals compared to naïve. **(C)** Synaptic gene ontology analysis of biological process considering DEGs in the vHip of stressed animals compared to naïve. Regulation of synaptic vesicle (SV) cycle was enriched in vHip of adolescent animals. * p < 0.05. **(D)** Normalized read counts of DEGs predicted to be involved in gene ontologies associated to synapse were significantly different among groups: *Vamp1* (t_14_ = 3.42, p = 0.004), *Exoc8* (t_14_ = 3.49, p = 0.003), *Picalm* (t_14_ = 3.96, p = 0.001), *Pheta1* (t_14_ = 3.49, p = 0.004), *Dtnbp1* (t_14_ = 3.49, p = 0.003) and *Syngr1* (t_14_ = 2.41, p = 0.03). **(E)** Normalized read counts of DEGs predicted to be involved in gene ontologies associated to redox system were significantly different among groups: *Gstm4* (t_14_ = 3.34, p = 0.005), *Gstt4* (t_14_ = 2.90, p = 0.01), *Tmem161a* (t_14_ = 2.75, p = 0.02), *Park7* (t_14_ = 2.97, p = 0.01) and *Rack1* (t_14_ = 2.21, p = 0.04). Data are shown as mean ± SEM. *p < 0.05; unpaired t-test.

To decipher the contribution of DEGs upon adolescent stress in the vHip, we next focused on adolescent stress-induced changes in the synaptic and redox system-associated pathways. Specifically, all synaptic-related genes were up-regulated (i.e., *Vamp1*, *Exoc8*, *Picalm*, *Pheta1,* and *Dtnbp1*), except for *Syngr1*, which was down-regulated in the vHip of adolescent stressed animals (Figure 2D). Redox system-related genes were all up-regulated, including those enriched in “Glutathione Transferase Activity” (*Gstm4* and *Gstt4*) and “Negative Regulation Of Oxidative-Stress Induced Neuron Death” (*Tmem161a* and *Park7*) pathways in the vHip of adolescent-stressed animals (Figure 2E). Moreover, *Rack1* gene was down-regulated and predicted to be involved in pathways related to “Negative Regulation Of Oxidative-Stress Induced Neuron Death”, “Negative Regulation Of Intrinsic Apoptotic Signaling Pathway In Response To DNA Damage” and “BH3 Domain Binding”.

### Adolescent stress leads to redox dysregulation and changes in mitochondrial respiratory function in the vHip

The transcriptional changes in vHip associated with oxidative stress and antioxidant response indicated a dysregulation in the redox system in adolescent stress animals. To dynamically characterize stress impact on redox homeostasis, we evaluated ROS levels one (PND41) and ten (PND51) days after adolescent stress. We found decreased levels of mitochondria-specific superoxide anion at PND51 without a difference at PND41 (Figure 3A). In contrast, adolescent stress increased hydrogen peroxide levels at PND41, with no changes at PND51 (Figure 3B). In conjunction, stressed animals at PND41 showed increased peroxidase activity without changes at PND 51 (Figure 3C). These results indicate a redox dysregulation caused by changes in free radical levels and enzymes that catalyze ROS.

**Figure 3.**
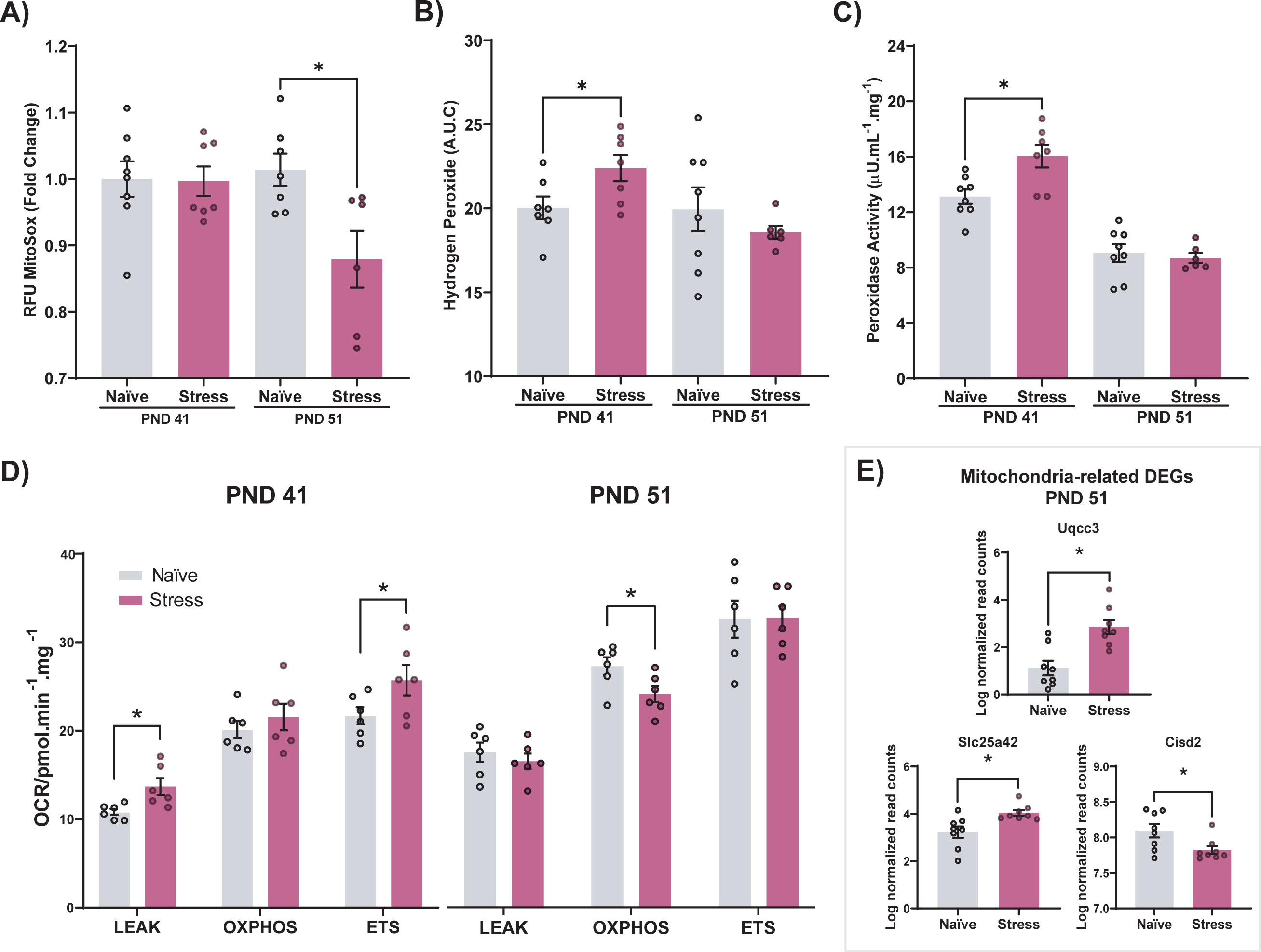
Adolescent stress leads to mitochondrial respiratory dysfunction and redox dysregulation. **(A)** Stress did not alter the mitochondrial superoxide anion levels at PND41 (t_13_ = 0.09, p = 0.93) but decreased at PND51 (t_12_ = 2.57, p = 0.02). RFU = Relative fluorescent units. **(B)** At PND41, hydrogen peroxide production was increased (t_13_ = 2.32, p = 0.04), with no changes at PND51 (t_12_ = 0.86, p = 0.40). A.U.C = area under curve. **(C)** Stress during adolescence increased peroxidase activity at PND41 (t_13_ = 3.10, p = 0.008), without changing it at PND51 (t_12_ = 0.44, p = 0.67) (n = 6–8/group). **(D)** Mitochondrial respiration in the vHip of naïve and stressed animals was evaluated on PND41 (n = 5–6/group) and PND51 (n = 6/group). Adolescent stress increased inducible protocol leak (t_10_ = 2.91, p = 0.02) and electron transport system capacity (ETS) (t_10_ = 2.84, p = 0.03). without changes in OXPHOS capacity (t_10_ = 0.83, p = 0.42). At PND51, stress decreased OXPHOS capacity (t_10_ = 2.34, p = 0.04) and did not impact Leak (t_10_ = 0.59, p = 0.56) or ETS (t_10_ = 0.12, p = 0.91). **(E)** Levels of gene expression from DEGs predicted to be associated with mitochondria are changed in the vHip adolescent stressed animals at PND51: *Uqcc3* (t_14_ = 4.04, p = 0.001), *Slc25a42* (t_14_ = 3.05, p = 0.009) and *Cisd2* (t_14_ = 2.47, p = 0.03). Data are shown as mean ± SEM. *p < 0.05; unpaired t-test.

The mitochondrial electron transport chain is the primary source of ROS, generating approximately 80% of cellular ROS in pathological conditions^28^, and its dysregulation can lead to an imbalance in physiological ROS concentration to produce oxidative stress^14^. Performing a high-resolution respirometry of the vHip samples at PND41, adolescent stressed animals showed increased respiration at LEAK and ETS states (Figure 3D), indicating increased mitochondrial uncoupling and electron transfer capacity, respectively; while at PND51, stressed animals showed decreased oxygen consumption at the OXPHOS state (Figure 3D), indicating reduced mitochondrial OXPHOS capacity in the vHip. Moreover, we also found several mitochondria-related DEGs in vHip of stressed animals at PND51 (*Uqcc3* and *Slc25a42* were down-regulated, and *Cisd2* was up-regulated; Figure 3E), reinforcing the impact of adolescent stress on vHip mitochondria-related genes and respiratory function.

### Adolescent stress enhanced GSSG in the vHip and serum

GSH is the principal endogenous antioxidant that maintains the brain’s redox balance and is a key regulator of the redox state of PV interneurons^13,29^. Through a reaction catalyzed by glutathione peroxidases, GSH donates its hydrogen to the highly reactive ROS and converts them into more stable compounds. In such reactions, two GSH molecules dimerize through a disulfide bond to form oxidized glutathione (GSSG). The GSH:GSSG ratio is a commonly used sensitive early biomarker of the whole-body redox status and a marker for oxidative stress^30^. In the vHip, stress did not change GSH (Figure 4A) but increase GSSG levels (Figure 4B) Also, GSH:GSSG ratio tends to decrease in the vHip of stressed animals (Figure 4C). In contrast, the serum of stressed animals showed high levels of GSH and GSSG (Figure 4D and E) and no change in GSH:GSSG ratio (Figure 4F) 10 days after stress. Then, we performed a Pearson correlation analysis to better understand the association between loss of sociability and cognitive impairment (Supplementary Figure 1A and B) with GSH and GSSG levels in the vHip and serum (Figure 4G). A negative correlation was observed between SIT and vHIP GSSG levels, suggesting that animals with loss of sociability showed higher levels of GSSG. SIT and NOR index were also negatively associated with serum GSH and GSSG levels. Additionally, vHIP GSSG levels positively correlate with serum GSSG.

**Figure 4.**
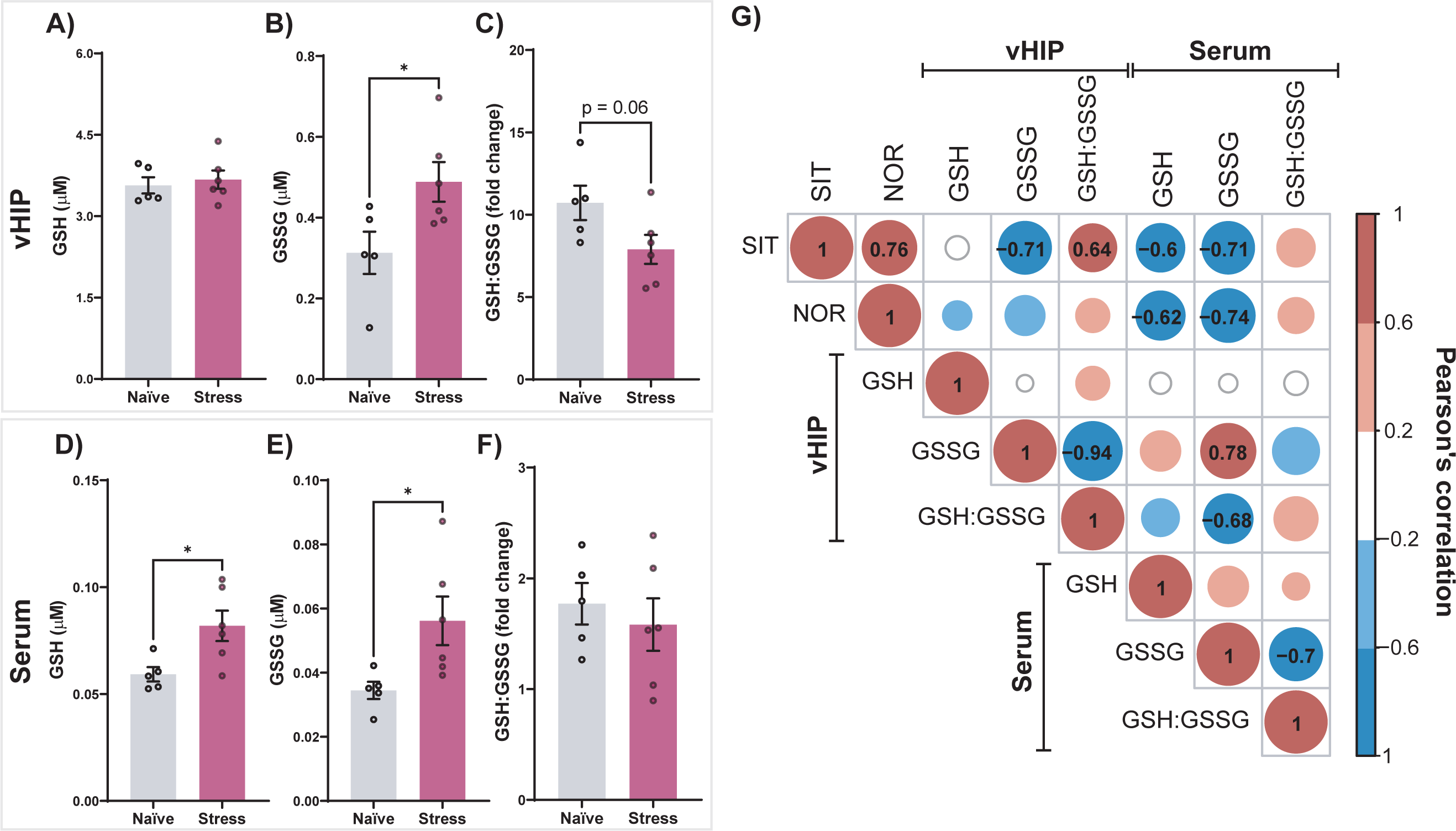
Effects of adolescent stress on GSH and GSSG in the vHip and serum. **(A)** While stress did not change GSH levels in the vHip (t_9_ = 0.46, p = 0.66), **(B)** GSSG form was increased (t_9_ = 2.44, p = 0.04), and **(C)** GSH:GSSG ratio tends to decrease (t_9_ = 2.08, p = 0.067). In the serum, stressed animals showed increased levels of **(D)** GSH (t_9_ = 2.69, p = 0.03) and **(E)** GSSG (t_9_ = 2.49, p = 0.04), **(F)** whereas no changes were observed in the GSH:GSSG ratio (t_9_ = 0.60, p = 0.56; n = 5–6/group). Data are shown as mean ± SEM. *p < 0.05; unpaired t-test. **(I)** Negative correlations were observed between SIT and GSSG (vHip and serum); SIT and GSH (serum); NOR and GSH (serum); and NOR and GSSG (serum); GSSG (vHip) and GSH:GSSG ratio (vHip); GSSG (serum) and GSH:GSSG ratio (serum); and GSSG (serum) and GSH:GSSG ratio (vHip). Positive correlations were observed between SIT and GSH:GSSG ratio (vHip); and GSSG (vHip) and GSSG (serum). r values are indicated in the matrix when correlations were p-adj < 0.05. Pearson’s correlation analysis. SIT = social interaction test; NOR = novel object recognition; GSH = reduced glutathione; GSSG = oxidized glutathione.

### Adolescent stress enhances vHip oxidative stress and its co-localization with PV interneurons

To assess potential cell DNA damage by adolescent stress, we performed immunohistochemistry for 8-OxodG, the most common DNA lesions resulting from ROS^31^, at PND 51, following behavioral tests. As expected, stressed animals showed more 8-OxodG fluorescence intensity in the vHip (Figure 5A), indicating DNA oxidative damage. PV interneurons are energy-demanding to support high-frequency neuronal synchronization^32^, and thus, they are vulnerable to redox dysregulation and more prone to oxidative damage after insults^7,33^. Given that adolescent stress decreased the number of PV-positive cells, we considered PV intensity fluorescence of the region of interest and characterized the degree of overlap with 8-OxodG fluorescence intensity. In addition, PV and 8-OxodG co-localization in the vHip increased after adolescent stress (Figure 5B), indicating that PV interneuron deficits caused by adolescent stress were accompanied by oxidative stress. Correlation analysis revealed that high levels of 8-OxodG were negatively associated with PV-positive cell number (Figure 5C). Also, a better behavioral performance in SIT negatively correlated with 8-OxodG intensity (Figure 5D). Such correlations were not observed for NOR (Figure 5E). Overall, our findings indicate that PV interneuron deficits and behaviors related to SCZ caused by adolescent stress were accompanied by oxidative stress.

**Figure 5.**
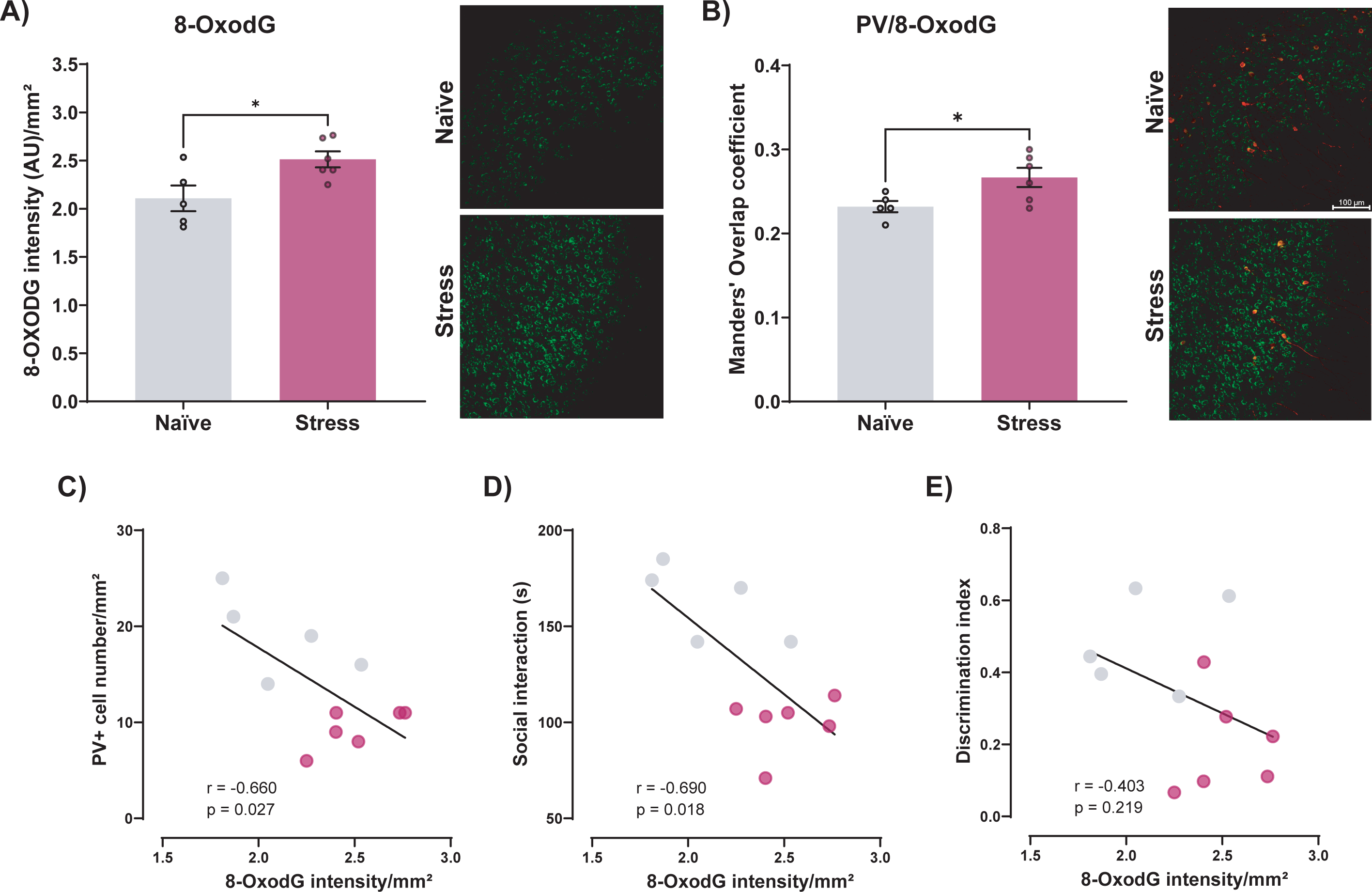
Oxidative damage in vHip PV interneurons after adolescent stress. **(A)** Stress increased the fluorescence intensity of the cell DNA damage marker 8-OxodG (t_9_ = 2.67, p = 0.02) and **(B)** its co-localization in PV-positive cells/mm² at PND51 (t_9_ = 2.47, p = 0.03). n = 5–6/group. AU = arbitrary units. Data are shown as mean ± SEM. *p < 0.05; unpaired t-test. **(C)** PV interneuron numbers/mm² were negatively correlated with 8-OxodG. **(D)** A negative correlation was observed between SIT and 8-OxodG **(E)** but was not significant for NOR and 8-OxodG association. r values are described in each graph after Pearson’s correlation analysis. *p < 0.05. SIT = social interaction test; NOR = novel object recognition.

## Discussion

Advancing our knowledge on markers and causative mechanisms that change adolescent behavior can contribute to discovering new strategies to prevent the emergence of neurodevelopmental disorders, such as SCZ. Here, using rats, we showed that: (1) adolescent stress led to a loss of sociability and cognitive impairment, reduction in PV-positive cells, an electrophysiological profile associated with SCZ, and transcriptional changes in the synaptic and redox system-related pathways; (2) adolescent stress-induced dysregulation in vHip ROS levels, possibly by impacting mitochondrial respiratory function; (3) high levels of GSSG were found in the vHip and serum of adolescent stressed animals, indicating a long-lasting antioxidant stress response; and, (4) adolescent-stress caused oxidative damage in PV interneurons and other cell types. Overall, our findings unveil new redox system-based insights and strengthen the role of mitochondria in developing brain abnormalities related to SCZ.

Stress during critical neurodevelopmental periods may lead to a glutamatergic overdrive onto PV interneurons, potentially changing the balance of E/I in cortical and limbic circuits that support multiple behaviors^34–36^. Common molecular features of numerous animal models to study SCZ include a reduced number of PV interneurons^37^. However, our previous meta-analysis revealed that only a few studies analyzed the impact of adolescent stress on the hippocampal PV interneurons^37^. Abnormal PV interneuron function is thought to lead to vHip hyperactivity, resulting in a hyperdopaminergic state^38^, a condition consistent with clinical observations in SCZ^39,40^. Like our previous studies^10^, adolescent stress caused hyperactivity of pyramidal neurons and diminished levels of PV interneurons and their PNNs in the vHip. Among synapse-related DEGs, the *Syngr* and *Dtnbp1* genes (associated with presynaptic vesicles in neuronal cells) are located on chromosome 22q13.1. and 6p22.3, respectively, both of which are recurrent linkage regions for SCZ^41,42^. Similar to stressed animals, *Syngr* expression is reduced in the postmortem brain of patients with SCZ^43^. While the *Dtnbp1* gene was up-regulated upon adolescent stress, its expression is decreased in patients with SCZ carrying a *Dtnbp1* risk haplotype, which was negatively associated with their cognitive abilities^44^.

Emerging evidence proposes that mitochondria play a prominent role in coordinating stress responses, making them a possible biological link between environmental insults and psychiatric outcomes^45^. Superoxide anion is the primary ROS generated by the electron transport chain and is detrimental to the overall viability of the cells. Once generated, superoxide anion gets rapidly converted by superoxide dismutase enzymes to hydrogen peroxide, the major contributor to cellular oxidative damage. Here, we found elevated hydrogen peroxide levels one day after stress, whereas superoxide anion levels were like control conditions. Indeed, at PND41, stressed rats exhibited an increased mitochondrial leak and electron transfer capacity. Proton leak and ROS generated from the electron transport chain are linked to each other since increased ROS activates mechanisms that promote proton leak, and in turn, increased proton leak reduces ROS production as a feedback loop^46^. Along with enhanced proton leak in adolescent stressed animals, the peroxidase activity was also elevated at PND41, revealing an antioxidant system activation to counteract the excessive presence of hydrogen peroxide levels immediately after stress exposure.

It is estimated that 80% of the total brain energy spending is accounted for processes related to neuronal signaling, highlighting the mitochondria-coupled respiration role in sustaining brain function^47^. The OXPHOS respiration in the vHip decreased in stressed animals at PND51, along with lower superoxide levels. Accordingly, exposure of animals to stress from PND 28–42 decreased the mitochondrial respiratory capacity of the nucleus accumbens during adulthood, which was associated with loss of sociability^48^. Notably, mitochondria-related DEGs in vHip included an up-regulation in ubiquinol-cytochrome C reductase complex assembly factor 3 (*Uqcc3*) and mitochondrial coenzyme A transporter (*Slc25a42*) genes. The Uqcc3 factor plays an important role in ATP production by stabilizing complex III proteins that transfer electrons to ubiquinol to cytochrome C^49^. Correspondingly, Slc25a42 protein exhibited a high transport affinity for CoA, ADP, and adenosine 3′,5′-diphosphate, in which its main function is to catalyze the entrance of CoA into the mitochondria in exchange for adenine nucleotides^50^. Although the upregulation of those genes suggests compensation for the reduced OXPHOS respiration, further causal experiments are required. In stressed animals, we also found down-regulation of the *Cisd2* gene, encoding for CDGSH iron sulfur domain 2. Cisd2 is localized on the endoplasmic reticulum and mitochondria-associated membranes, being crucial in the regulation of cytosolic Ca^2+^ homeostasis and mitochondrial function^51^. For instance, *Cisd2* deficiency in mice causes mitochondrial fission and dysfunction accompanied by autophagic cell death, along with behavioral phenotype suggestive of premature aging^52^.

GSH is the most prominent antioxidant in the brain since it maintains the homeostasis of redox states in cells and prevents oxidative damage in the tissue^53^. Supporting the transcriptomic changes in the “glutathione transferase activity” pathway, in which DEGs predicted to be involved in glutathione metabolism were up-regulated (i.e., *Gstm4* and *Gstt4*), we found that GSSG was increased in the vHip of stressed animals at PND51, pointing to a previous increase of ROS after adolescent stress. Indeed, GSH:GSSG ratio, a marker for oxidative stress, tended to decrease ten days after stress. In contrast to our results, a significant increase in GSH was found in the hippocampus of rats exposed to GBR 12909, a dopamine reuptake inhibitor, during early postnatal development (PND 5–16)^54^. Also, exposing mice to 8 weeks of post-weaning social isolation decreased the cortical and striatal GSH:GSSG ratio in rats, suggesting a reduction in GSSG and/or an increase in GSH in these brain regions^55^. Notably, a transgenic mouse model of impaired synthesis of GSH (*Gclm* KO mice) showed a 70% reduction in brain GSH levels and exhibit behavioral homologies with SCZ^56,57^. The genetically compromised GSH synthesis affects PV interneurons in the vHip but not the dorsal hippocampus, which alters emotion-related behaviors^33^. Additionally, in *Gclm* KO mice, PV interneurons and PNNs deficits emerged in a spatiotemporal sequence that paralleled regional anterior cingulate cortex maturation, suggesting that GSH deficit delays the maturation of PV interneurons and their PNNs, which correlated with oxidative DNA damage brain regions throughout development^7,58^. Here, we also found that GSH and GSSG increased in the serum of stressed animals, indicating a possible compensatory/protective mechanism by enhancing the peripheral antioxidant system^55,59^. In patients with psychosis who reported early traumatic experiences, higher blood glutathione peroxidase levels, a peripheral marker reflecting low brain GSH levels, were described and associated with more severe clinical symptoms^60^.

Although many studies have observed an association between oxidative stress and environmental risk factors for mental disorders^61^, limited evidence indicates that oxidative stress is prevalent in the PV interneuron population when evaluated in the prefrontal cortex and dentate gyrus after early-life or juvenile insults^62,63^. Here, we found that animals exposed to stress during adolescence showed vHip higher levels of oxidative stress markers at PND51, which was co-localized with PV. Additionally, our transcriptomic analysis points to “Negative Regulation Of Oxidative Stress-Induced Neuron Death” and pathway, including up-regulation of *Tmem161a* (Transmembrane Protein 161A or Adaptive Response To Oxidative Stress Protein 29 – Aro-29) and *Park7* (Parkinsonism associated deglycase or DJ-1), and down-regulation of *Rack1* (Receptor for activated C Kinase 1) genes. While the mechanism by which Tmem161a/Aro-29 reduced levels of oxidant-induced DNA damage and apoptosis remains unclear^64^, Park7/DJ-1 acts as an oxidative stress sensor by promoting expression of several Nrf2 target genes, including those associated with glutathione metabolism^65^. Furthermore, Park7/DJ-1 interacts with Rack1, protecting cortical neurons from hydrogen peroxide-induced apoptosis in vitro and in vivo models^66,67^.

Our findings indicate a dynamic and temporal change in ROS levels and mitochondrial respiratory function, leading to oxidative stress on PV interneurons and, consequently, deficits in the vHip E/I balance and behavioral abnormalities. Under chronic stress, an additional energetic burden is required to support allostasis and stress-induced energy needs^68–70^. An allostatic overload occurs when mediators responsible for helping stress adaptation become harmful, i.e., the energetic costs of stress limit energy to sustain the demands of proper mental health and supersede the available energetic reserve. Allostatic overload manifests at multiple levels of biological systems (organism, cellular, molecular), such as accumulation of damage and accelerated aging. This perspective would reconcile the long-lasting mitochondrial and redox changes with reductions in PV-positive cells in the vHip of stressed animals. Indeed, the bioenergetic cost of adolescent stress seems too high to be sustainable by PV interneurons. Notably, the energetic view of stress and health brings potential mechanisms of stress resilience, since increasing the threshold for allostatic overload can improve the ability of the organism to tolerate chronic stress. Our study gathers novel information highlighting important targeted strategies that ameliorate brain energy levels, such as enhancing the antioxidant system response or mitochondrial respiratory function, to prevent and/or rescue the E/I imbalance and the maladaptive stress overload on adolescent behavioral outcomes.

## Acknowledgments

The authors acknowledge Sabrina Baroni’s technical support in performing the RNA integrity analysis. The authors thank Marco Antonio de Carvalho, Eleni Tamburus Gomes, and Eliane Aparecida Antunes Maciel for technical assistance.

## Data availability

The data that support the findings of this study and the R scripts analyses are available from the corresponding author upon reasonable request.

