## Supplemental material for "Adolescent stress-induced ventral hippocampus redox dysregulation underlies behavioral deficits and excitatory/inhibitory imbalance related to schizophrenia"

### *Supplemental Information*

### EXTENDED METHODS AND MATERIALS

#### 1. Animals

Male and female Sprague-Dawley rats (postnatal day, PND 70) were obtained from the University of São Paulo Central Animal Facility, Ribeirão Preto. Rats were allowed to acclimatize for one week in the animal facility before breeding. Mating was confirmed by the presence of spermatozoa in the vaginal smear, and birthday defined PND 0. At PND 21, pups were weaned. We opted to use only the male offspring, as we previously found that female adolescent rats did not show behavioral and electrophysiological changes induced by the stress protocol used here[1]. Animals were randomly assigned to all experimental groups, each cage devoted to a specific experimental procedure. Rats were housed (2 – 3 animals per cage) in a temperature (22 °C) and humidity (47%) controlled environment (12 h light/dark cycle; lights on at 6 AM) with water and food ad libitum. All procedures were approved by the Ribeirão Preto Medical School Ethics Committee (155/2018 and 248/2019), which follows Brazilian and International regulations.

#### 2. Stress procedure

Animals were exposed to inescapable footshock (FS; from PND 31 – 40) daily and three restraint stress (RS) sessions (PD31, 32, and 40). Using this protocol, we previously found marked behavioral and electrophysiological changes that persist until adulthood[2–4]. Briefly, rats were exposed to one session of FS per day for 10 consecutive days. In each session, animals were placed in a Plexiglas chamber with a grid floor of 0.48 cm stainless steel rods spaced 1.6 cm apart (EP107R, Insight, Brazil). Twenty-five FS (1.0 mA, 2 s) were delivered pseudo-randomly (5 cycles of 30, 60, 40, 60, and 90 s). Immediately after receiving FS, rats were submitted to RS for 1 h in Plexiglas cylindrical size-adjusted restraint tubes on the first, second, and last day of FS exposure. Cylinders measured 14.0 × 3.9 cm (length × diameter) for rats at

PND 31 – 32 and 20.3 × 5.1 cm for rats at PND 40, ventilated by holes (1 cm diameter). Naïve animals were left undisturbed in their home cages.

### 3. Behavioral tests

#### 3.1. Social Interaction Test (SIT)

Animals were placed in the center of a circular arena (60 cm height × 65 cm diameter) and allowed to explore it for 5 min. Then, an unfamiliar male Sprague-Dawley rat (PND 50 – 54) was introduced into the arena as a social target for 10 min. The social interaction time was measured when the tested animal was sniffing the unfamiliar rat's anogenital region, head, or body and when they were following, crawling over and under each other. A blind experimenter quantified the social interaction time.

#### 3.2. Novel Object Recognition (NOR)

Two hours after the SIT, each animal was habituated in a circular arena (60 cm height × 65 cm diameter) for 10 min. NOR test was conducted in the same circular arena 24 h later. Animals were submitted to two trials separated by 1 h. During the first trial (acquisition trial, T1), rats were placed in the arena containing two identical objects for 5 min. For the second trial (retention trial, T2), one of the objects presented in T1 was replaced by an unknown (novel) object. Animals were then placed back in the arena for 5 min. Object exploration was defined as when the animal faced the object at 2 cm of distance or less while watching, licking, sniffing, or touching it with the forepaws while sniffing. A blind experimenter quantified object exploration. Recognition memory was assessed using the discrimination index:

$$[(t_{novel\ object} - t_{familiar\ object})/(t_{novel\ object} + t_{familiar\ object})]$$

### 4. In vivo recordings of vHip pyramidal neurons

Rats were anesthetized with chloral hydrate (400 mg/kg, i.p.) and mounted on a stereotaxic frame. The coordinates for the vHip were 5.3 mm posterior from bregma, 4.4 mm lateral to the midline, and 5.5 – 8.0 mm ventral from the brain surface. Electrodes were lowered through 6 tracks inside the vHip. Pyramidal neurons were identified by typical electrophysiological characteristics such as firing rate (average up to 2 Hz) and action potential shape (half–width> 0.4 ms)[2,4]. The firing rate of these neurons was measured. Each identified pyramidal neuron was recorded for 1–3 min. After the electrophysiological recordings, the electrode sites were marked via iontophoretic ejection of Chicago Sky Blue dye from the electrode (20 μA constant negative current, 20 min). Then, rats were euthanized by an overdose of chloral hydrate; the brains were removed, fixed for at least 24 h in 8% paraformaldehyde (PFA), cryoprotected in 30% sucrose, and sectioned for histological confirmation of the electrode sites.

### 5. Gene expression profiling from the rat PFC

#### 5.1. RNA isolation

Following the behavioral tests on PND 51, animals were anesthetized (25% urethane, 1 mL/100g/rat) and perfused with cold 0.01M phosphate-buffered saline (PBS, pH=7.4). vHip from both hemispheres was collected and snapped frozen in liquid nitrogen until use for RNA extraction (n=8/group). To this end, we used RNAqueous-Micro Total RNA Isolation Kit (ThermoFisher Scientific; #AM1931), according to the manufacturer's instructions.

#### 5.2. Bulk RNA-sequencing

The extracted RNA was used for performing the transcriptomic analysis from the vHip of naïve and adolescent-stressed rats using bulk RNA barcoding and sequencing, as previously described[5,6]. Briefly, the RNA samples were reverse transcribed with individual barcoded oligodT primers. Then, all samples were pooled together, and the second strand synthesis generated the double-stranded cDNA via the nick translation method. Illumina-compatible libraries were prepared by tag-mentation of 5 ng of full-length double-stranded cDNA. Then, the library was amplified, profiled, and sequenced using the Illumina NextSeq 500 platform.

#### 5.3. Transcriptomic analysis

Following a quality assessment with FastQC[7], gene reads were mapped with HISAT2 onto Rnor_6.0/rn6 genome assembly for Rattus norvegicus[8]. Mapped reads were counted for each gene locus using the featureCounts function of the subread (2.0.2) package. We normalized count data by size factor and applied a variance stabilizing transformation for visualization purposes[9]. Low-abundance genes were removed before data normalization, keeping only genes with at least ten reads in all samples. Subsequently, we performed a generalized linear model to assess differentially expressed genes (DEGs) using the DESeq2 package[10]. p-values were corrected for multiple testing using the Benjamini–Hochberg method[11]. Transcriptomic analysis was performed in R (R Core Team, 2014).

#### 5.4. Gene set enrichment analysis

Only DEGs with a p-value $< 0.01$ were explored for enriched gene sets and function. Gene set enrichment analyses were derived from bioinformatics resource systems (web servers) for functional annotation and enrichment analyses of gene lists. Enrichr was used for gene ontology enrichments since it integrates knowledge from many high-profile projects to provide synthesized information about mammalian genes and gene sets[12–14]. Also, Synaptic Gene Ontologies (SynGo) annotations were used to examine gene set enrichment associated to synaptic process. Mitochondrial-related DEGs were retrieved from MitoCarta3.0, an inventory of mammalian mitochondrial proteins and pathways[15].

### 6. Biomolecular analyses

#### 6.1. High-resolution respirometry

High-resolution respirometry is an approach to assess cellular oxygen consumption by evaluating mitochondrial respiratory states and maximal mitochondrial electron transport system capacity in fresh-permeabilized tissues[16]. Naïve and stressed rats (n = 6/group) were anesthetized (25% urethane, 1 mL/100 g/rat), and 2 mg of fresh vHip samples from both hemispheres were finely cut into pieces and used in the experiments. The experiments were made in duplicate, as previously described[17]. The samples were permeabilized in BIOPS solution (2.7 mM EGTA, 20 mM imidazole, 20 mM taurine, 50 mM acid 2-(N-morfolino) ethanesulfonic potassium, 0.5 mM dithiothreitol, 6.5 mM MgCl2, 15 mM phosphocreatine, 0.57 mM ATP, pH 7.1) containing 0.01% saponin for 5 min at 4ºoC, then carefully transferred to the chambers of an Oxygraph-2k respirometer (Oroboros, Austria), containing 2.1 mL of air saturated respiration medium MIR05 (0.5 mM EGTA, 3 mM MgCl2, 60 mM K-lactobionate, 20 mM taurine, 10 mM KH2PO4, 20 mM HEPES, 110 mM sucrose, 1 g/L albumin, pH 7.1). Basal respiratory rates were determined after adding 9 mM glutamate and 5 mM malate. OXPHOS rates were measured after adding 1 mM ADP to the chambers. This was followed by the addition of oligomycin (1 μg/mL), an inhibitor of the ATP synthase, to measure respiration not linked to ATP production (inducible proton leak). To uncouple respiration and measure maximal capacity rates of the Electron Transport Chain (ETC) 2 pulses of the mitochondrial uncoupler carbonyl cyanide m-chlorophenylhydrazone (CCCP, 1 μM) were added. Finally, residual oxygen consumption (Rox) rates were determined by adding 1 mM NaCN. Rox rates were subtracted from all other measurements.

### 6.2. MitoSOX™ and AmplexRed® assays

An independent experimental group was employed to measure the release of hydrogen peroxide from vHip samples and the superoxide production by mitochondria. For these purposes, we used Amplex® Red Hydrogen Peroxide/Peroxidase Assay Kit (ThermoFisher, A22188) and MitoSOX™ mitochondrial superoxide indicator (ThermoFisher, M36008) assay[18]. Naïve and adolescent stressed animals (n = 6–8/group) were anesthetized with 25% urethane (1 mL/100 g/rat), perfused with ice-cold 0.01 M PBS, and decapitated. vHip tissues were immediately collected, cut into approximately 1 $mm^3$ cubes, and added into HAM – F12 culture medium containing B27 supplement without antioxidants and collagenase type IV (0.05%). The tissues were incubated under gentle agitation for 30 min at 37 °C for cell dissociation. Next, a cell suspension was obtained by pipetting up and down the tissue, followed by centrifugation at 1200 g for 10 min. The collagenase-containing medium was discarded, and the pellet was resuspended in HBSS to a 200 µg/µL dilution.

MitoSOX™ assay was used to quantify mitochondrial superoxide anion in the vHip. It corresponds to a DHE derivative, conjugated with a long aliphatic chain containing a phosphonium group, which targets the probe to the mitochondrial matrix. The MitoSOX™ reacts with the superoxide anion, forming an intensely fluorescent compound. The fluorescence intensities were determined spectrophotometrically in the FlexStation apparatus (Molecular Devices, USA) according to the manufacturer's guidelines. The values were expressed for t = 0 as RFU.

The amount of hydrogen peroxide ($H_2O_2$) was measured using the Amplex® Red Assay Kit according to the manufacturer's instructions. Amplex® Red (AR) reagent is a highly sensitive and stable probe for detecting $H_2O_2$ in living samples, especially for cell-based assays. It is based on the following reaction, conducted in the presence of horseradish peroxidase (HRP):

$$AR + H_2O_2 \xrightarrow{\text{HRP}} Resorufin + O_2$$

Resorufin (7-Hydroxy-3H-phenoxazin-3-one) is a highly fluorescent product with an excitation maximum at 530 – 570 nm and an emission maximum at 580 – 590 nm, allowing to monitor $H_2O_2$ extracellular flux fluorometrically, and ROS production by living cells. Fluorescence intensity was evaluated every 5 min for 20 min in the FlexStation apparatus (Molecular Devices, USA). Values were expressed as $H_2O_2$/µmol.min-1.mg-1. Then, we performed the area under curve measurement to report the hydrogen peroxide production in the intact cells during 20 min of incubation.

Next, we performed the tissue peroxidase activity assay. The HRP enzyme, which catalyzes the reaction of $H_2O_2$ with the AmplexRed® assay, was removed from the reaction buffer. Then, an excess of $H_2O_2$ was added to each sample, allowing the evaluation of the endogenous peroxidase activity of the cell suspensions. The values were evaluated for t = 0 and expressed as enzyme activity per reaction volume (µU.mL-1.mg-1).

#### 6.3. Glutathione/Glutathione disulfide (GSH/GSSG) assay

vHip GSH and GSSG levels were measured by a green fluorescence assay kit (ab205811, Abcam, UK). For that end, at PND 51, animals were anesthetized (urethane 25%, 1 mL/100 g/rat) and perfused with cold 0.01 M PBS. vHip from both hemispheres was collected and snapped frozen in liquid nitrogen until use for GSH total extraction (n = 6/group). 20 mg of tissue were resuspended in 400 µL of ice-cold Mammalian Lysis Buffer and homogeneized with 10-15 passes. Then, samples were centrifuged at 12,000 x g for 10 min at 4 °C. Supernatants were collected and deproteinized by adding 1 volume of trichloroacetic acid (TCA, 100% w/a) into 5 volumes of samples. After 10 min of incubation, samples were centrifuged at 12,000 x g for 5 min at 4 °C. Samples were neutralized by adding sodium bicarbonate ($NaHCO_3$) to the supernatant and vortex. After confirming the pH was between 5 – 6, samples were centrifuged

(13,000 x g for 15 min at 4 °C), and the supernatant was used to measure reduced GSH and total GSH. The fluorescence intensities of the samples were determined spectrophotometrically in the FlexStation apparatus (Molecular Devices, USA) according to the manufacturer's guidelines.

The following linear regression obtained changes in fluorescence intensity with GSH concentration:

$$Log(_y) = (A + B) * Log\ (x)$$

Then, the dilution factor corrected the final concentration (μM) of GSH and/or Total GSH + GSSG. The concentration of GSSG (μM) in the test samples was calculated as the following equation:

$$GSSG = {}^{total\ GSH - GSH}/_{2}$$

### 7. Immunofluorescence

All immunohistochemical procedures followed previously reported protocol with minor modifications[4]. In brief, rats were deeply anesthetized with urethane 25% (1 mL/100 g/rat) and transcardially perfused with 0.01 M PBS, followed by 4% PFA in 0.01 M PBS (pH = 7.6). Then, brains were removed, post-fixed in 4% PFA for 2 h, and stored in 30% sucrose. Serial 30 μm-thick coronal sections of the vHip were collected using a cryostat (CM-1900, Leica). For each animal, five to six sections 300 μm apart spanning the rostrocaudal axis of the vHip (within −4.8 mm to −6.3 mm from bregma) were collected and stained. Specifically, sections were incubated in a combination of 1% normal goat serum, 0.1% Triton X-100, rabbit anti-PV antibody (1:2000, Swant, PV 25), biotinylated Wisteria floribunda agglutinin (WFA; 1:1000 dilution, Vector Labs, #B1355) and mouse Anti-8-OxodG antibody (1:500, Abcam, ab62623) for 24 h at 4 °C. The sections were then incubated with a mixture of 1% normal goat serum, goat anti-rabbit Alexa

Fluor 488 (1:1000, Abcam, ab150077), Alexa Fluor 594 conjugated to streptavidin (1:1000, Abcam, ab272189), and goat anti-mouse Alexa Fluor 647 (1:1000, Abcam, ab150115) for 90 min. The sections were mounted with Fluoroshield Mounting Medium with DAPI (Abcam, ab104139) to visualize the border of the vSub.

#### 7.1. Image acquisition and analysis

For image acquisition, the focus was set on PV-positive cells for imaging, and digital images were obtained using Leica Application Suite X (Leica Microsystems). Under 20x magnification, the vSub regions of each rostrocaudal section were imaged by z-stacks (512 × 512 μm-images along the medial-temporal axis) using a confocal microscope (SP5, Leica). An abrupt widening of the pyramidal layer defined the border between CA1 and vSub. The boundary between the subiculum and presubiculum was characterized by a sharp reduction in PV intensity and a decrease in cell size visualized by DAPI. For cell count, only the pyramidal cell layers, where most PV interneurons were located, were counted. The exposure time for PV was calibrated such that most of the PV-positive cells in the naïve group were visible and within the dynamic range, and all subsequent images of the remaining age groups were taken at an identical exposure. Similar techniques were applied to PNN, 8-OxodG, and DAPI to identify optimal exposure time. PNNs were identified by staining for WFA, a lectin that selectively labels residues of glycoproteins within the PNNs. While this does not show the PNN structure explicitly, nor are PNNs located exclusively around PV neurons[19], counterstaining for PV enables us to tell if the structure is indeed a perisomatic PNN encompassing PV interneurons.

For analysis, acquired images were first converted to maximum projection (z-stacks). The PV positive cell count was then performed using Fiji software, cell counts plugin. Also, the fluorescence intensity of PNN and 8-OxodG labeling were analyzed in Fiji software. Data were reported as integrated density (arbitrary units, AU).PV positive cell number and fluorescence intensity AU were normalized by total area ($mm^2$). For co-localization analysis, the intensity of a pixel in one channel was evaluated against the corresponding pixel in the second channel of a dual-color image. The Manders' overlap coefficients were obtained using the co-localization threshold plugin in the Fiji software to measure the amount of PV/PNN and PV/8-OxodG co-localization.

### 8. Statistical analyses

All data were subjected to tests to verify the homogeneity of variances (Bartlett's test) and if they followed a normal distribution (Shapiro–Wilk test). Those that met these parameters were subjected to parametric analysis (Student's t-test, one-way or two-way ANOVA followed by post-hoc Turkey's test), and data were expressed as mean ± SEM. Otherwise, they were subjected to nonparametric analysis (Mann-Whitney test), and data were represented as boxes and whiskers (minimum and maximum value). Significant differences were indicated by $p < 0.05$. Statistical analyses were performed with Prism 8.0 (Graphpad Software Inc.) and R (R Core Team, 2014).

17. Santos JR Dos, Bortolanza M, Ferrari GD, et al. One-Week High-Intensity Interval Training Increases Hippocampal Plasticity and Mitochondrial Content without Changes

in Redox State. *Antioxidants 2020, Vol 9, Page 445*. 2020;9(5):445. doi:10.3390/ANTIOX9050445

18. Votyakova T V., Reynolds IJ. ΔΨm-Dependent and -independent production of reactive oxygen species by rat brain mitochondria. *J Neurochem*. 2001;79(2):266-277. doi:10.1046/J.1471-4159.2001.00548.X

19. Lensjø KK, Christensen AC, Tennøe S, Fyhn M, Hafting T. Differential Expression and Cell-Type Specificity of Perineuronal Nets in Hippocampus, Medial Entorhinal Cortex, and Visual Cortex Examined in the Rat and Mouse. *eNeuro*. 2017;4(3):379-395. doi:10.1523/ENEURO.0379-16.2017
