## Supplementary figures and images for "Adolescent stress-induced ventral hippocampus redox dysregulation underlies behavioral deficits and excitatory/inhibitory imbalance related to schizophrenia"

### Supplemental Figure 1

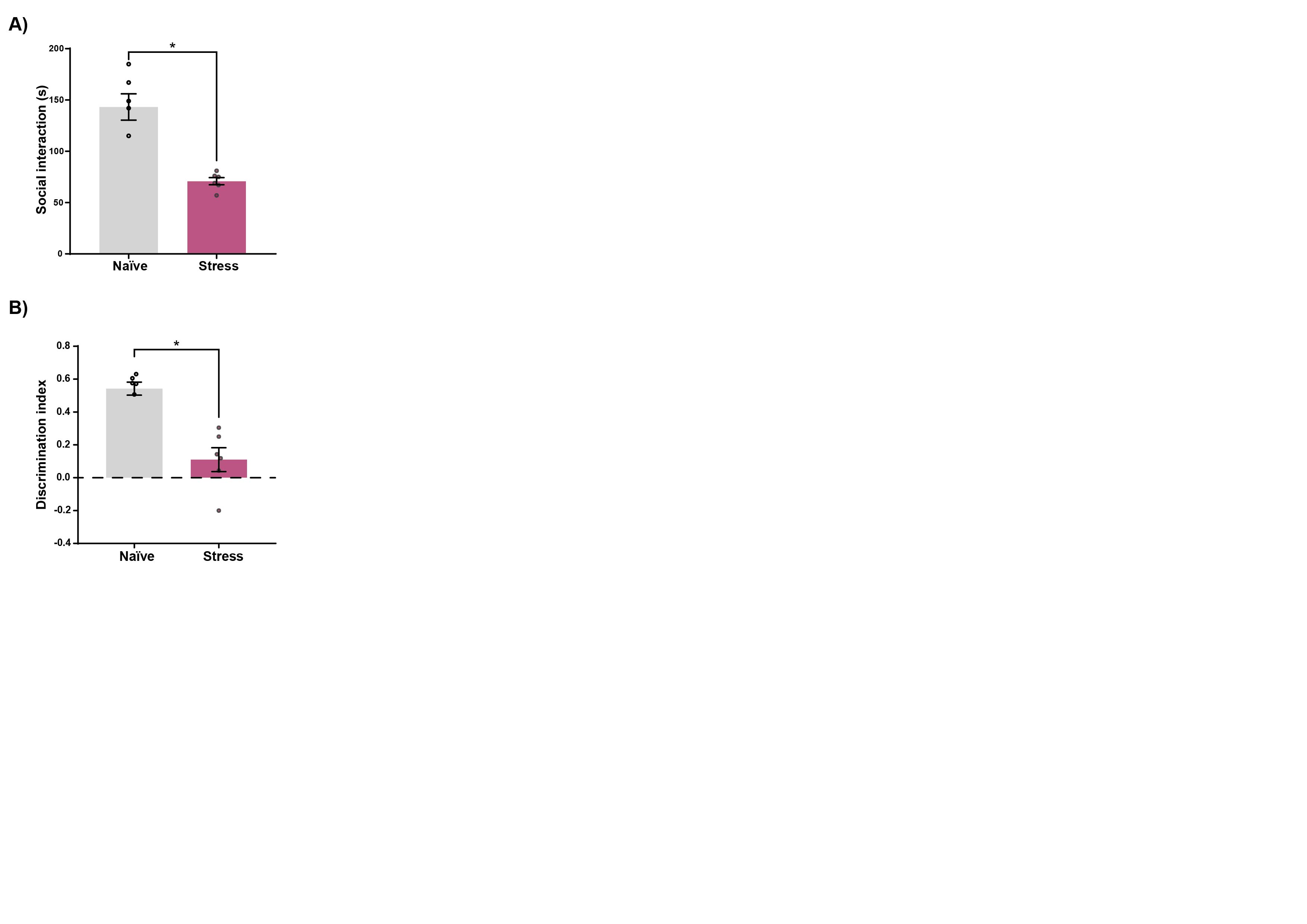
